# Substrate preferences, phylogenetic and biochemical properties of proteolytic bacteria present in the digestive tract of Nile tilapia

**DOI:** 10.1101/2021.10.24.465423

**Authors:** Tanim Jabid Hossain, Mukta Das, Ferdausi Ali, Sumaiya Islam Chowdhury, Subrina Akter Zedny

**Affiliations:** Department of Biochemistry and Molecular Biology, University of Chittagong, Chattogram 4331, Bangladesh; Biochemistry and Pathogenesis of Microbes Research Group, Chattogram 4331, Bangladesh; Department of Microbiology, University of Chittagong, Chattogram 4331, Bangladesh

**Keywords:** Nile tilapia, gut bacteria, bacterial extracellular protease, proteolytic activity, 16S rRNA gene based phylogeny, substrate preference

## Abstract

Vertebrate intestine appears an excellent source of proteolytic bacteria for industrial and probiotic use. We therefore aimed obtaining the gut-associated proteolytic species of Nile tilapia. We’ve isolated twenty six bacterial strains from its intestinal tract, seven of which showed exoprotease activity with the formation of clear halos on skim milk. Their depolymerization ability was further assessed on three distinct proteins including casein, gelatin and albumin. All the isolates could successfully hydrolyze the three substrates indicating relatively broad specificity of their secreted proteases. Molecular taxonomy and phylogeny of the proteolytic isolates were determined based on their 16S rRNA gene barcoding which suggested that the seven strains belong to three phyla viz. Firmicutes, Proteobacteria and Actinobacteria, distributed across the genera *Priestia, Citrobacter, Pseudomonas, Stenotrophomonas, Burkholderia, Providencia* and *Micrococcus*. The isolates were further characterized by a comprehensive study of their morphological, cultural, cellular and biochemical properties which were consistent with the phylogenetic annotations. To reveal their proteolytic capacity alongside substrate preferences, enzyme-production was determined by the diffusion assay. The *Pseudomonas, Stenotrophomonas* and *Micrococcus* isolates appeared most promising with maximum protease production on casein, gelatin and albumin media respectively. Our findings present valuable insights into the phylogenetic and biochemical properties of gut-associated proteolytic strains of Nile tilapia.

## Introduction

Proteolytic enzymes, also called proteases, catalyze degradation of proteins and peptides by hydrolytic cleavage of peptide bonds [1]. Being essential for cell growth and differentiation, the proteolytic enzymes are ubiquitous in biological systems [2]. Microorganisms produce a vast diversity of intracellular and extracellular proteases. While the intracellular proteases play essential functions in cellular biochemistry, physiology, and regulatory aspects, the extracellular proteases provide carbon and nitrogen sources to cells by degrading extracellular proteins into small peptides and amino acids that can be transported into the cells [3]. Aside from their importance from biological point of view, the proteolytic activity is sought in numerous industrial processes, for example, in the detergent, leather, fabric and food industries, in pharmacology and drug manufacture, waste management, animal feed preparations etc. [4, 5]. Furthermore, proteases are commonly used as basic research tools in many biochemical investigations. For example, in protein identification, unknown proteins are subjected to trypsin digestion into small peptides for their subsequent analysis by mass spectrometry [6]. Other important applications in research include peptide synthesis, peptide sequencing, digestion of unwanted proteins in purified samples as in the DNA and RNA purifications, Klenow fragment production and so on [7–10]. With the total annual sales of about 1.5–1.8 billion USD, proteases, therefore, account for about 60% of the global enzyme sales constituting the largest product-segment of industrial enzymes [11]. Although the proteolytic enzymes can be obtained from many of the organisms, those derived from microbes especially bacteria are preferred for the large-scale production since bacteria are the easiest, cheapest and fastest to grow in a relatively small and simple set-up and are also suitable to genetic manipulation for increased production. Microbial proteases were also found more active and stable at extreme conditions than those of the plant or animal origin [12]. Therefore, the microbial enzymes can be obtained in abundant quantities on a regular basis and with a uniform quality [13]. Hence, many commercially important enzymes including proteases are generally obtained from a variety of bacterial species.

Recently, use of the protease producing bacteria is gaining increasing acceptance in aquaculture industry, world’s fastest growing food production sector [14]. The proteolytic bacteria if included in aquaculture may serve multiple purposes such as (1) improved digestion of protein-rich substances present in the host’s natural diet and in commercial feed resulting in an increased growth of the host [15]; (2) enhancement of nonspecific immune response in the host [16]; (3) reduction of organic pollutants produced in aquaculture from the undigested feed [17] etc. Besides, as compared to exogenous proteases, use of the protease producing microbes are more ecofriendly and easy in the application in aquaculture [18].

Nile tilapia is the third most important aquaculture species by volume having an enormous economic value [19]. For its high popularity among consumers and its easy and inexpensive method of farming, tilapia has become the most widely cultivated fish worldwide [20, 21]. The fish has a versatile eating habit and consumes phytoplankton, zooplanktons, macrophytes, insects, detritus, nematodes etc. in its diet [22]. Being a herbivorous-omnivorous fish without a true stomach, and with phytoplankton and plant debris comprising a major portion of its diet, Nile tilapia generally lacks pepsin and the role of pepsin is taken over by alkaline proteases which are more active in an alkaline environment [23]. Supplementing its feed with bacteria secreting extracellular proteases, therefore, appears highly beneficial to its cultivation.

To address the increasing demand of protease producing bacteria in industry, research and aquaculture, we focused on obtaining proteolytic strains from the gastrointestinal tract (GIT) of Nile tilapia. Fish GIT has been recognized as an excellent source of bacteria producing extracellular hydrolytic enzymes [15], and there is also a general consensus that the bacteria to be included in the animal feed should be isolated from GIT of the animals where they will be applied [18]. Consequently, we’ve isolated cultivable GI bacteria from Nile tilapia and screened them for protease production. The positive isolates were all identified and extensively characterized based on their genetic and biochemical properties and sugar fermentation abilities. Moreover, their substrate preferences as well as depolymerization capacities on various protein substrates were also studied.

## Materials and Methods

### Preparation of intestinal sample

For isolation of bacteria, intestinal sample was prepared from fish purchased from a local market. Entire digestive tract of the fish was removed by aseptic surgery and its external surface was thoroughly washed with autoclaved distilled water and then sterilized using 70% v/v ethanol. Internal contents of the digestive tract were then squeezed out and collected in a beaker. Inside of the digestive tract was then rinsed well with sterile water which was also added to the internal contents.

### Isolation of bacteria

Bacteria present in the intestinal sample were isolated as previously described with minor modifications [15]. Briefly, 100 μl of the intestinal sample and its 10-fold serial dilutions (10^0^ through 10^-6^) were spread on nutrient agar (NA; 5 g/L peptone, 3 g/L yeast extract, 5 g/L NaCl, 18 g/L agar; pH 7) and Luria-Bertani (LB) agar (10 g/L tryptone, 5 g/L yeast extract, 10 g/L NaCl, 18 g/L agar; pH 7) [24] plates and incubated at 30°C for 24 to 48 h. All morphologically distinct colonies were selected and streaked on fresh NA and LB agar plates to obtain pure cultures [25].

### Preparation of culture stock

Cells from the colony of pure culture was inoculated to nutrient and LB broth and incubated at 37°C. After 24 h of growth, 500 μL of the culture was transferred to a cryo-vial, sterile glycerol was added to the final concentration of 20% v/v and preserved at −80°C for further analysis.

### Culture conditions

The isolates were routinely maintained in LB media at 30°C, unless otherwise noted. Each isolate was revived from its glycerol stock by transferring cells to 2 to 5 ml LB broth by a sterile loop and grown overnight in an orbital shaker at 180 rpm at 30°C. 1% v/v of this activated overnight culture was transferred to 10 ml fresh broth, incubated at 30°C for 24 h and used for subsequent analysis.

### Screening for proteolytic activity

To detect presence of extracellular proteolytic activity, 10 μl of a 0.8 OD_600_ culture of each isolate was spot-inoculated on the surface of skim milk agar media (5 g/L peptone, 2.5 g/L yeast extract, 1 g/L dextrose, 28 g/L skim milk powder, 18 g/L agar; pH 7) as well as NA and LB media each supplemented with 1% (w/v) skim milk powder and incubated at 30°C for 24 to 48 h. Isolates that showed zones of clear halo surrounding the colonies were considered positive for protease production.

### Morphological, cultural and biochemical characterization

Determination of morphological, cultural and biochemical properties of the isolates and their fermentation of various carbohydrates were carried out by methods described previously [25, 26].

### 16S rRNA gene amplification and sequencing

16S rRNA gene of each isolate was amplified from its genomic DNA using GoTaq G2 Hot Start Master Mix (Promega) and the purified PCR products were sequenced using BigDye Terminator v3.1 Cycle Sequencing Kit according to a previous report [15].

### Sequence deposition

The 16S rRNA genes sequenced in the present study have been deposited in the GenBank database under the accession numbers OK287066 to OK287072.

### Taxonomic analysis

Taxonomic annotation of the proteolytic isolates was carried out by analysis of their 16S rRNA gene sequences with nucleotide BLAST of NCBI [27], RDP classifier and seqmatch [28] and Silva ACT: Alignment, Classification and Tree Service [29]. All parameters were set to default values with the only exception in BLAST search where the “Max target sequences” was set to 1000. Phylotypes in the BLAST searches were determined by considering the query coverage, percent identity, maximum and total scores, and the total number of hits obtained for the query sequence against a particular genus or species. Organisms with an ambiguous taxonomic description such as enrichment culture clones, uncultured bacteria or unclassified bacteria were not taken into consideration [31]. NCBI taxonomy browser was followed to obtained taxonomic hierarchy of the isolates [32].

### Phylogenetic analysis

Phylogenetic analysis of the isolates was performed essentially as previously described [33]. 16S rRNA gene sequence of the isolates, and 700 bp of their nearest type strains, and the top hit strains in BLAST results were aligned by Muscle or ClustalW algorithms in Molecular Evolutionary Genetics Analysis (MEGA) software, version X [34]. The closest type strain for each isolate was found by using EzBioCloud’s 16S-based ID [35], and their sequences were collected from the EzBioCloud database having the accession numbers CP001628, LASD01000006, FLYB01000015, JJMH01000057, HQ888847, BAMA01000316, LDJN01000038. Two additional strains used in the alignment for each isolate were selected from the top hits in BLAST search results and their sequences were obtained from GenBank database with the accession numbers MW198159.1, MT509874.1, MT509997.1, MK033338.1, MN420979.1, MH341969.1, MT533939.1, MT033093.1, MK571729.1, MK640708.1, KY913809.1, EU307934.1, MT555731.1, MT649753.1. A phylogenetic tree of the aligned sequences was built by maximum likelihood (ML) method using Tamura-Nei algorithm with 100 bootstrap replications and by neighbor joining (NJ) method using Maximum Composite Likelihood algorithm in MEGA as described in [31].

### Determination of substrate specificity

Ability of the proteolytic isolates to hydrolyze casein, gelatin and bovine serum albumin (BSA) was examined based on the formation of clear halos around colonies streaked on NA and LB media supplemented with 1% (w/v) of each substrate as described above.

### Estimation of relative activity

To determine relative proteolytic activity, the isolates were grown on media containing 1% (w/v) of casein, gelatin or BSA at 30°C for 48 h. The diameter of the zone of clearance and that of the colonies were measured. Relative activity (RA) was then calculated using the formula, RA□=□(colony diameter□+□halo zone diameter)/colony diameter [15].

### Statistical analysis

All experiments were performed at least three times separately, averaged and the standard deviation was generated. The data were presented as the mean ± standard deviation displayed as error bars.

## Results

### Proteolytic activity of the gut associated bacteria

In this study, we aim to isolate and characterize proteolytic strains in the gut flora of nilotica. To this end, 26 of its gut associated culture-dependent strains were isolated and screened for their ability to produce extracellular proteases on skim milk agar plates. Only 7 of the isolates (designated as TGB1 to TGB7) showed proteolytic activity as indicated by the formation of clear halos on media due to the depolymerization of casein in skim milk (supplementary Figure S1a). To further evaluate their proteolytic aptitude, enzyme activity was assessed on three different protein substrates including casein, gelatin and BSA. All the seven isolates were found capable of degrading the three substrates which indicate relatively broad specificity of their secreted proteases.

### Taxonomic and phylogenetic characteristics

Molecular taxonomy of the protease producing strains was determined by homology and phylogeny analysis of their 16S rRNA gene sequences to those in various databases. The sequences were subjected to a battery of 16S rRNA gene based methods for their identification. Results of the sequence analysis and subsequent phylotype assignments are presented in Table 1. Nucleotide blast of the sequences against those in GenBank and EzBioCloud databases showed a high sequence-similarity, with the percent identities higher than 99% to the respective sequences of *Priestia, Citrobacter, Pseudomonas, Stenotrophomonas, Burkholderia, Providencia* and *Micrococcus* (Table 1). The taxonomic assignment was also supported by other classification platforms such as RDP classifier, EzBioCloud 16S-based ID and Silva ACT (Table 1) confirming the taxonomic annotations to at least genus level. Phylotypes of the isolates each belonging to a separate genus indicates a very high diversity among the isolates without a single genus found predominant over the others. Considering their phylotypes along the taxonomic hierarchy, it was observed that the isolates belong to the phyla Firmicutes, Proteobacteria and Actinobacteria, with Proteobacteria being highly dominant (~72%).

**Table 1.**
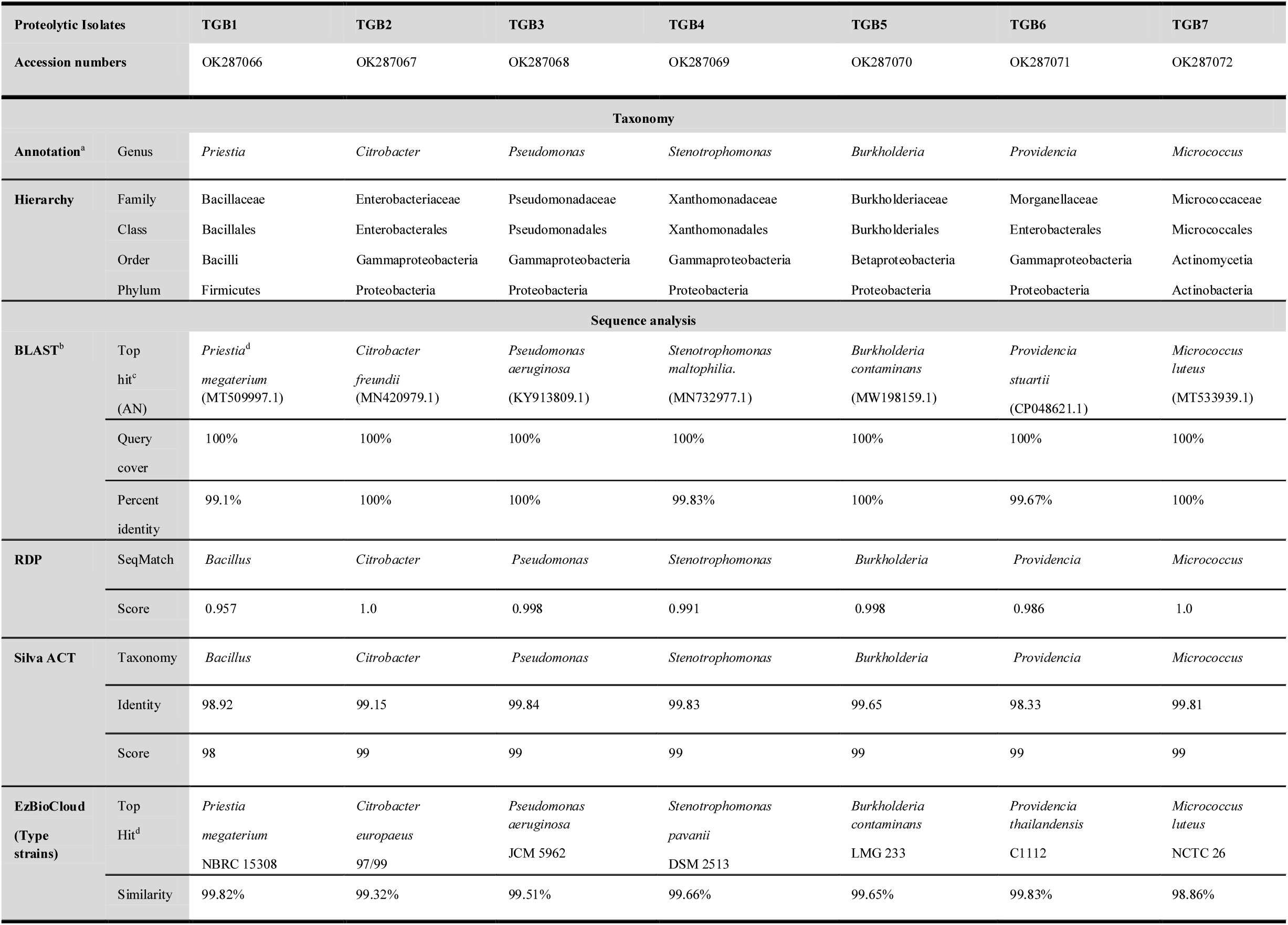
Taxonomic affiliations of the isolates based on analysis of their 16S rRNA gene sequences.

Phylogenetic tree based on homology of the 16S rRNA genes of the isolates with their closest GenBank strains and type strains is depicted in Figure 1. The phylogenetic analysis showed a clear congruence with taxonomic assignments of the isolates. Each isolate formed a separate cluster with its nearest type strain and GenBank strains of the same species, located at similar distances.

**Figure 1.**
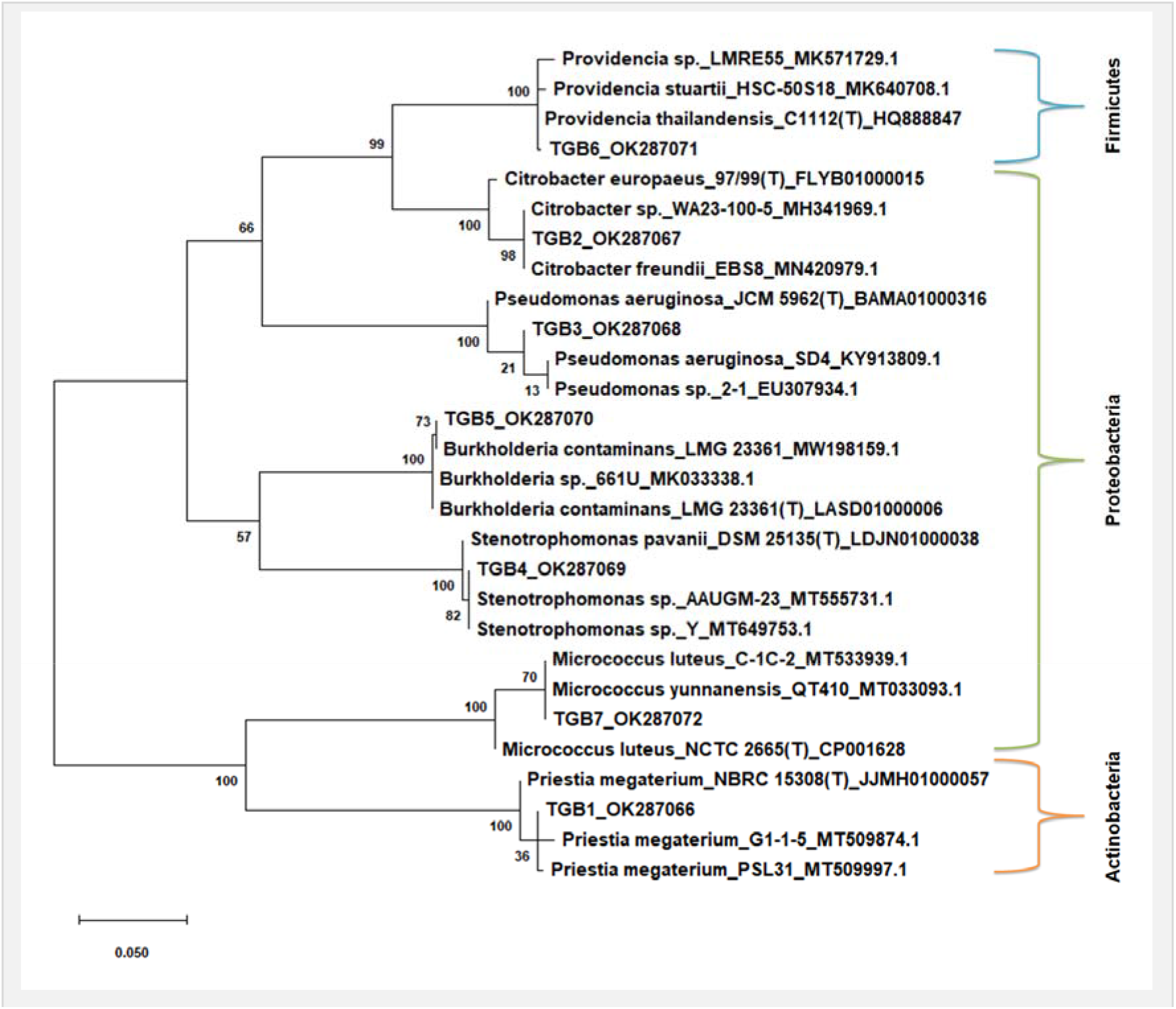
Phylogenetic orthogonal tree depicting distribution and relationships in the protease producing isolates and their closest type strains and GenBank strains. Species names are followed by strain names and accession numbers separated by underscores. Type strains are indicated by (T). Tree was built by ML (shown here) and NJ methods both producing the same results. Tree inference was performed as described in “Materials and Methods”.

### Morphological, cellular and biochemical properties

Morphological, cultural and cellular characteristics of the proteolytic isolates and their biochemical properties are summarized in Tables 2, 3 and 4. Cell morphology showed that most of the isolates were Gram negative rods although TGB1 and TGB7 appeared Gram positive and TGB7 was found to be a coccus (Table 3). All isolates could produce catalase and most of them also produced H_2_S. In MR-VP tests, however, all isolates showed negative results. Extracellular amylase activity was detected in three of the isolates including the *Priestia* (TGB1), *Citrobacter* (TGB2) and *Stenotrophomonas* (TGB4) strains.

**Table 2.** Morphological and cultural characteristics of the protease producing strains.

| **Isolates** | **Colony on**<br>**NA medium** | **Colony**<br>**color** | **Appearance in**<br>**broth culture** | **Oxygen**<br>**requirement** |
| --- | --- | --- | --- | --- |
| **TGB1** | Irregular, raised with undulate edge | Dull white | Turbidity with pellicle and sediment in the bottom of the tube | Aerobic |
| **TGB2** | Irregular, raised with undulate edge | Dull white | Turbidity with pellicle and sediment in the bottom of the tube | Facultative anaerobe |
| **TGB3** | Circular, entire, low convex with regular edge | Yellowish white | Dense turbidity and sediment in the bottom of the tube. | Facultative anaerobe |
| **TGB4** | Circular, raised with regular edge | Dull white | Uniform turbidity | Aerobic |
| **TGB5** | Punctiform, flat with regular edge on Nutrient Medium | Dull white | Turbidity with pellicle and sediment in the bottom of the tube | Facultative anaerobe |
| **TGB6** | Punctiform, convex with regular edge | Yellowish | Uniform Turbidity and sediment in the bottom of the tube. | Facultative anaerobe |
| **TGB7** | Circular, raised with regular edge | Yellowish | Uniform Turbidity and sediment in the bottom of the tube. | Facultative anaerobe |

**Table 3.** Cellular characteristics of the isolates.

| **Isolates** | **Cell shape** | **Cellular arrangement** | **Motility** | **Gram staining** |
| --- | --- | --- | --- | --- |
| **TGB1** | Straight rod | Single or pairs | Motile | Gram positive |
| **TGB2** | Straight rod | Single | Motile | Gram negative |
| **TGB3** | Straight/slightly curved rod | Single | Motile | Gram negative |
| **TGB4** | Straight rod | Single | Motile | Gram negative |
| **TGB5** | Rod | Single | Non-motile | Gram negative |
| **TGB6** | Straight rod | Single | Non-motile | Gram negative |
| **TGB7** | Cocci | Tetrads/pairs | Non-motile | Gram positive |

**Table 4.** Biochemical properties and sugar fermentation of the protease producing strains.

| Isolates | TGB1 | TGB2 | TGB3 | TGB4 | TGB5 | TGB6 | TGB7 |
| --- | --- | --- | --- | --- | --- | --- | --- |
| Basic biochemical properties |  |  |  |  |  |  |  |
| Catalase | + | + | + | + | + | + | + |
| Oxidase | + | - | + | - | - | - | + |
| Indole | - | - | - | + | - | - | - |
| H<sub>2</sub>S | + | + | - | + | + | - | Weekly + |
| MR | - | - | - | - | - | Weekly + | - |
| VP | - | - | - | - | - | - | - |
| Starch hydrolysis | + | + | - | + | - | - | - |
| Sugar fermentation |  |  |  |  |  |  |  |
| Arabinose | - | + | - | - | - | - | - |
| Glucose | + | + | + | + | + | - | - |
| Fructose | - | - | - | + | + | - | - |
| Galactose | + | - | - | - | + | - | + |
| Sucrose | - | - | - | + | + | - | - |
| Starch | + | - | - | - | - | - | - |
| Mannitol | - | + | + | - | - | - | + |
| Raffinose | - | - | - | - | - | - | - |

+ = positive result, - = negative result

Fermentation tests with carbohydrates including various mono, di, tri and polysaccharides showed that the isolates had a rather limited metabolic capacity. Glucose was the sugar fermented by most (5/7) isolates. A maximum of five sugars could be fermented by the *Burkholderia* (TGB5) isolate. Overall, the cultural and biochemical properties of the isolates largely complied to their phylogenetic affiliations as described in the Bergey’s manual of systematic bacteriology [36].

### Protease producing capacity and substrate preferences

To evaluate protease producing capacity of the isolates on different substrates, a general estimate of their protease production was performed based on diffusion of the secreted proteases across culture medium and presented as relative activity (RA) [18] in Figure 2. Three different isolates, *Pseudomonas* (TGB3), *Stenotrophomonas* (TGB4) and *Micrococcus* (TGB7), were found producing the maximum amount of protease on the casein, gelatin and albumin media respectively. A relatively higher production on casein media was also exhibited by the *Providencia* (TGB6) and *Micrococcus* (TGB7) isolates, and on gelatin media by the *Pseudomonas* (TGB3), *Burkholderia* (TGB5) and *Micrococcus* (TGB7) isolates. The *Micrococcus* (TGB7) strain, therefore, appeared to be the only isolate efficient with any of the three substrates. In general, most of the isolates showed protease producing capacity in the order of gelatin > casein > BSA; exceptions were the *Pseudomonas* (TGB3) and *Providencia* (TGB6) isolates in which the order was casein > gelatin > BSA. Such a pattern suggests that protease released by the bacteria might have relatively higher preferences for casein and gelatin over albumin.

**Figure 2.**
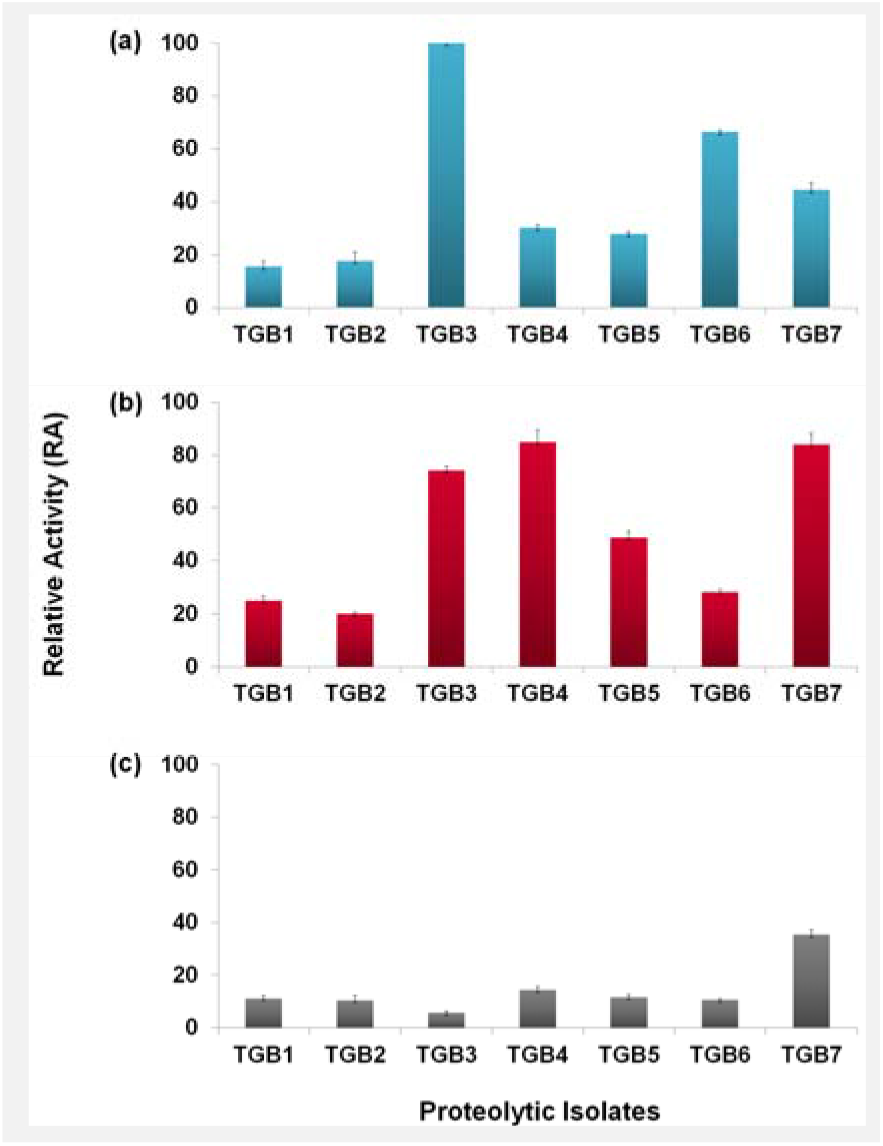
Protease producing capacity of the isolates on (a) casein, (b) gelatin and (c) albumin used as substrates in the medium, presented as relative activity (RA). Error bars represent standard deviation of the mean (n=3).

## Discussion

We carried out this study to obtain proteolytic bacteria from the GIT of Nile tilapia since bacteria producing extracellular protease are demonstrated to have the potential to be used as probiotic agents in fish feed; moreover they also comprise a valuable source of enzymes for research and industrial use. We’ve already discussed importance of the protease producing bacteria in research, aquaculture and industries in the introduction section. The beneficial gastrointestinal flora has been recognized in recent research as the most suitable candidates intended for probiotic use [37]. Accordingly, we’ve isolated and studied gut bacteria of tilapia and detected proteolytic activity in about 27% of the isolates. The fact that the major fraction (73%) of the isolates lacked protease producing ability is not unusual considering that Nile tilapia has a herbivorous-omnivorous feeding habit. In our previous research on microbial hydrolytic enzymes, we found proteolytic activity in 50% of the intestinal bacteria in Bombay duck which, however, is a carnivore [15]. The diet of a carnivore is supposed to be rich in protein substances and largely devoid of plant based materials. As a result the proteolytic strains are expected to be dominant among the GI flora of a carnivorous fish. Consistent with this perception, Bairagi et al. reported relatively high densities of cellulolytic and amylolytic strains in tilapia although proteolytic isolates were also found in considerable numbers [38]. Similarly, Kar and Ghosh found higher populations of proteolytic bacteria in the carnivorous fish *Channa punctatus* than that in the herbivore *Labeo rohita* [39]. Although all these previous studies including ours arrived at the same conclusion suggesting it to be a general phenomenon, to fully confirm if it is indeed the case for Nile tilapia to have relatively lower proportion of proteolytic strains, an extensive study should be performed with large number of samples analyzed individually by both culture-dependent and metagenomic methods. But the primary objective of this work being obtaining proteolytic strains for downstream applications, it was outside of the scope.

The protease producing isolates of the present study were all identified genetically from their 16S rRNA gene analysis which was further supported from their morphological and biochemical properties. The isolates appeared taxonomically diverse at the genus level each belonging to a separate phylotype. Few of the similar phylotypes have been previously documented in the GIT of Nile tilapia. For example, species of *Bacillus* (*B. megaterium*; reclassified as *Priestia megaterium*), *Citrobacter*, and *Burkholderia* were commonly isolated from Nile tilapia [37, 40–43], and therefore seems to be autochthonous to this fish. Moreover, these three species recovered from tilapia intestine were reported to possess extracellular protease activity and other beneficial properties, and are being considered as probiotic candidates for Nile tilapia [37, 44, 45]. Although not frequently, but the other four genera identified in our analysis, *Pseudomonas, Stenotrophomonas, Micrococcus* and *Providencia*, have also been described among the intestinal bacteria of Nile tilapia [42, 46–48]. Whatever the source of their isolation is, species of all the seven genera were reported producing extracellular proteases [49–56]. At the phylum level, Proteobacteria were found dominant over the other two phyla, Firmicutes and Actinobacteria in our study. In general, the above three phyla have been commonly found among the gut flora of this fish. For example, similar to our findings, Wu et al. also identified species which belonged only to the above three phyla where species of Firmicutes were found more dominant in the gut of Nile tilapia fed with woody forages [42]. In a culture-independent study using metagenomic approach Bereded et al. reported that the gut microbiota of Nile tilapia were dominated by two more phyla Cyanobacteria, Fusobacteria in addition to the above three [57].

All isolates of this study showed ability to degrade three different proteins including casein, gelatin and albumin with different degrees of degradation efficiencies and substrate preferences. However, albumin turned out to be relatively less preferred. Most isolates showed higher affinity for gelatin followed by casein and albumin as also previously reported, for example, in proteases from a *Photobacterium* sp. and a *Brevibacillus brevis* isolate [58, 59]. Three of our isolates, on the other hand, showed highest preference for casein which has been commonly observed in previous studies as well [60–64].

In summary, we’ve isolated and identified protease producing bacteria in the gut of Nile tilapia. We revealed morphological, cellular and biochemical properties of the proteolytic isolates and showed that their secreted proteases could hydrolyze casein, gelatin and albumin with different depolymerization capacities. Further investigations on ability of the proteases to digest proteins in aquaculture feed, elucidation of their structural and catalytic properties for industrial exploitations, and occurrence of additional beneficial properties in the proteolytic isolates, are needed.

## Acknowledgments

We are grateful to Prof. Dr. Md. Monirul Islam, Dept. of Biochemistry and Molecular Biology, University of Chittagong (BMB, CU) for his kind support with laboratory equipment. We thank members of the Biochemistry and Pathogenesis of Microbes (BPM) Research Group, BMB, CU for their various help in this project.

## Author contributions

TJH conceived and designed the study; MD, FA, TJH and SIC performed experiments; SIC contributed to sequence study and TJH performed the phylogenetic analysis; TJH and FA interpreted the data; TJH wrote and prepared the manuscript; SAZ assisted in writing introduction; SAZ and FA helped in information collection; all authors approved the final manuscript.

## References

[1] Jakubke H-D, Kuhl P, Könnecke A. Basic principles of protease - catalyzed peptide bond formation. Angew Chem Int Ed Eng. 1985;24:85–93.

[2] Gupta R, Beg Q, Lorenz P. Bacterial alkaline proteases: molecular approaches and industrial applications. Appl Microbiol Biotechnol. 2002;59:15–32.

[3] Waschkowitz T, Rockstroh S, Daniel R. Isolation and characterization of metalloproteases with a novel domain structure by construction and screening of metagenomic libraries. Appl Environ Microbiol. 2009;75:2506–2516.

[4] Razzaq A, Shamsi S, Ali A, Ali Q, Sajjad M, Malik A, et al. Microbial proteases applications. Front Bioeng Biotechnol. 2019;7:110.

[5] Zhu D, Wu Q, Hua L. Industrial enzymes. In: Moo-Young M, Editor. Comprehensive Biotechnology. Oxford: Pergamon;2019. p. 1–13.

[6] Graves PR, Haystead TAJ. Molecular biologist’s guide to proteomics. Microbiol Mol Biol Rev. 2002;66:39–63.

[7] Białkowska AM, Morawski K, Florczak T. Extremophilic proteases as novel and efficient tools in short peptide synthesis. J Ind Microbiol Biotechnol. 2017;44:1325–1342.

[8] Yang H, Li Y-C, Zhao M-Z, Wu F-L, Wang X, Xiao W-D, et al. Precision de novo peptide sequencing using mirror proteases of Ac-LysargiNase and trypsin for large-scale proteomics. Mol Cell Proteomics. 2019;18:773–785.

[9] Theron LW, Divol B. Microbial aspartic proteases: current and potential applications in industry. Appl Microbiol Biotechnol. 2014;98:8853–8868.

[10] Eun H-M. DNA polymerases. In: Eun H-M, Editor. Enzymology Primer for Recombinant DNA Technology. San Diego: Academic Press;1996. p. 345–489.

[11] Olajuyigbe FM, Falade AM. Purification and partial characterization of serine alkaline metalloprotease from *Bacillus brevis* MWB-01. Bioresour Bioprocess. 2014;1:8.

[12] Cui H, Yang M, Wang L, Xian C-J. Identification of a new marine bacterial strain SD8 and optimization of its culture conditions for producing alkaline protease. Plos ONE. 2015;10:e0146067.

[13] Martínez-Medina GA, Barragán AP, Ruiz HA, Ilyina A, Martinez HJ, Rodríguez-Jasso R, et al. Fungal proteases and production of bioactive peptides for the food industry. In: Kuddus M, Editor. Enzymes in Food Biotechnology. Cambridge: Academic Press;2019. p. 221–246.

[14] Tacon AGJ. Trends in global aquaculture and aquafeed production: 2000–2017. Rev Fish Sci Aquacult. 2020;28:43–56.

[15] Hossain TJ, Chowdhury SI, Mozumder HA, Ali F, Chowdhury MNA, Rahman N, et al. Hydrolytic exoenzymes produced by bacteria isolated and identified from the gastrointestinal tract of bombay duck. Front Microbiol. 2020;11:02097.

[16] Selim KM, Reda RM. Improvement of immunity and disease resistance in the Nile tilapia, *Oreochromis niloticus*, by dietary supplementation with *Bacillus amyloliquefaciens*. Fish Shellfish Immunol. 2015;44:496–503.

[17] Su H, Xiao Z, Yu K, Huang Q, Wang G, Wang Y, et al. Diversity of cultivable protease-producing bacteria and their extracellular proteases associated to scleractinian corals. PeerJ. 2020;8:e9055.

[18] Amin M. Marine protease-producing bacterium and its potential use as an abalone probiont. Aquacult Rep. 2018;12:30–35.

[19] Maas RM, Deng Y, Dersjant-Li Y, Petit J, Verdegem MCJ, Sharma JW, Kokou F. Exogenous enzymes and probiotics alter digestion kinetics, volatile fatty acid content and microbial interactions in the gut of Nile tilapia. Sci Rep. 2021;11:8221.

[20] Anshary H, Kurniawan RA, Sriwulan S, Ramli R, Baxa D. Isolation and molecular identification of the etiological agents of streptococcosis in Nile tilapia (*Oreochromis niloticus*) cultured in net cages in Lake Sentani, Papua, Indonesia. SpringerPlus. 2014;3:627.

[21] Champneys T, Castaldo G, Consuegra S, Garcia de Leaniz C. Density-dependent changes in neophobia and stress-coping styles in the world’s oldest farmed fish. Royal Soc Open Sci. 2018;5:181473.

[22] Njiru M, Okeyo-Owuor J, Muchiri M, Cowx I. Shifts in the food of Nile tilapia, *Oreochromis niloticus* (L.) in Lake Victoria, Kenya. African J Ecol. 2004;42:163–170.

[23] Moyle PB, Cech JJ. Fishes: An introduction to ichthyology. New Jersey: Pearson Prentice Hall;2004.

[24] Guzman MLCD, Arcega KSG, Cabigao J-MNR, Su GLS. Isolation and identification of heavy metal-tolerant bacteria from an industrial site as a possible source for bioremediation of cadmium, lead, and nickel. Adv Environ Biol. 2016;10:10–16.

[25] Hossain TJ, Alam MK, Sikdar D. Chemical and microbiological quality assessment of raw and processed liquid market milks of Bangladesh. Cont J Food Sci Technol. 2011;5:6–17.

[26] Carter GR. Diagnostic Procedure in Veterinary Bacteriology and Mycology. Amsterdam: Elsevier;1990.

[27] Zhang Z, Schwartz S, Wagner L, Milner W. A greedy algorithm for aligning DNA sequences. J Comput Biol. 2000;7:203–214.

[28] Wang Q, Garrity GM, Tiedje JM, Cole J. Naive Bayesian classifier for rapid assignment of rRNA sequences into the new bacterial taxonomy. Appl Environ Microbiol. 2007;73:5261–5267.

[29] Pruesse E, Peplies J, Glöckner FO. SINA: Accurate high-throughput multiple sequence alignment of ribosomal RNA genes. Bioinf. 2012;28:1823–1829.

[30] Larsen MV, Cosentino S, Lukjancenko O, Saputa D, Rasmussen S, Hasman H, et al. Benchmarking of methods for genomic taxonomy. J Clin Microbiol. 2014;52:1529–1539.

[31] Hossain TJ, Manabe S, Ito Y, Iida T, Kosono S, Ueda K, et al. Enrichment and characterization of a bacterial mixture capable of utilizing *C*-mannosyl tryptophan as a carbon source. Glycoconjugate J. 2018;35:165–176.

[32] Schoch CL, Ciufo S, Domrachev M, Hotton CL, Kannan S, Khovanskaya R, et al. NCBI Taxonomy: a comprehensive update on curation, resources and tools. Database (Oxford) 2020. 2020:baaa062.

[33] Ferdausi A, Sharup D, Hossain TJ, Chowdhury SI, Zedny SA, Das T, et al. Production optimization, stability, and oil emulsifying potential of biosurfactants from selected bacteria isolated from oil contaminated sites. R Soc Open Science. 2021;DOI: 10.1098/rsos.211003.

[34] Kumar S, Stecher G, Li M, Knyaz C, Tamura K. MEGA X: Molecular evolutionary genetics analysis across computing platforms. Mol Biol Evol. 2018;35:1547–1549.

[35] Yoon S-H, Ha S-M, Kwon S, Lim J, Kim Y, Seo H, Chun J. Introducing EzBioCloud: a taxonomically united database of 16S rRNA gene sequences and whole-genome assemblies. Int J Syst Evol Microbiol. 2017;67:1613–1617.

[36] Garrity GM, Bell JA, Lilburn TG. Taxonomic Outline of the Prokaryotes: Bergey’s Manual of Systematic Bacteriology. New York: Springer-Verlag; 2004.

[37] Reda RM, Selim KM, El-Sayed HM, El-Hady MA. In Vitro selection and identification of potential probiotics isolated from the gastrointestinal tract of Nile tilapia, *Oreochromis niloticus*. Probiotics Antimicrob Proteins. 2018;10:692–703.

[38] Bairagi A, Ghosh KS, Sen SK, Ray AK. Enzyme producing bacterial flora isolated from fish digestive tracts. Aquacult Int. 2002;10:109–121.

[39] Kar N, Ghosh K. Enzyme Producing bacteria in the gastrointestinal tracts of *Labeo rohita* (Hamilton) and *Channa punctatus* (Bloch). Turk J Fish Aquat Sci. 2008;8:115–120.

[40] Molinari L, Scoaris D, Pedroso R, Bittencourt N, Nakamura C, Ueda-Nakamura T, et al. Bacterial microflora in the gastrointestinal tract of Nile tilapia, *Oreochromis niloticus*, cultured in a semi-intensive system. Acta Sci Biol Sci. 2003;25:267–271.

[41] Saha S, Roy RN, Sen SK, Ray AK. Characterization of cellulase-producing bacteria from the digestive tract of tilapia, *Oreochromis mossambica* (Peters) and grass carp, *Ctenopharyngodon idella* (Valenciennes). Aquacult Res. 2006;37:380–388.

[42] Wu F, Chen B, Liu S, Xia X, Gao L, Zhang X, Pan Q. Effects of woody forages on biodiversity and bioactivity of aerobic culturable gut bacteria of tilapia (*Oreochromis niloticus*). Plos ONE. 2020;15:e0235560.

[43] Haygood AM, Jha R. Strategies to modulate the intestinal microbiota of Tilapia (*Oreochromis* sp.) in aquaculture: a review. Rev Aquacult. 2018;10:320–333.

[44] Afrilasari W, Widanarni, Meryandini A. Effect of probiotic *Bacillus megaterium* PTB 1.4 on the population of intestinal microflora, digestive enzyme activity and the growth of catfish (Clarias sp.). HAYATI J Biosci. 2016;23:168–172.

[45] Zorriehzahra MJ, Delshad ST, Adel M, Tiwari R, Karthik K, Dhama K, et al. Probiotics as beneficial microbes in aquaculture: an update on their multiple modes of action: a review. Vet Q. 2016;36:228–241.

[46] Yang C, Jiang M, Lu X, Wen H. Effects of dietary protein level on the gut microbiome and nutrient metabolism in tilapia (*Oreochromis niloticus)*. Anim. 2021;11:1024.

[47] Zaky MMM, Ibrahim ME. Screening of bacterial and fungal biota associated with *Oreochromis niloticus* in lake manzala and its impact on human health. Health. 2017;9:697–714.

[48] Boari CA, Pereira GI, Valeriano C, Silva BS, Morais VM, Figueiredo HC, et al. Bacterial ecology of tilapia fresh fillets and some factors that can influence their microbial quality. Food Sci Technol. 2008;28:863–867.

[49] Biedendieck R, Knuuti T, Moore SJ, Jahn D. The “beauty in the beast”— the multiple uses of *Priestia megaterium* in biotechnology. Appl Microbiol Biotechnol. 2021;105:5719–5737.

[50] Nicodème M, Grill J-P, Humbert G, Gaillard J-L. Extracellular protease activity of different *Pseudomonas* strains: dependence of proteolytic activity on culture conditions. J Appl Microbiol. 2005;99:641–648.

[51] Asker MMS, Mahmoud MG, El Shebwy K, Abd el Aziz MS. Purification and characterization of two thermostable protease fractions from *Bacillus megaterium*. J Genet Eng Biotechnol. 2013;11:103–109.

[52] Ray AK, Roy T, Mondal S, Ringø E. Identification of gut-associated amylase, cellulase and protease-producing bacteria in three species of Indian major carps. Aquacult Res. 2010;41:1462–1469.

[53] Lee M-A, Liu Y. Sequencing and characterization of a novel serine metalloprotease from *Burkholderia pseudomallei*. FEMS Microbiol Lett. 2000;192:67–72.

[54] Miyaji T, Otta Y, Shibata T, Mitsui K, Nakagawa T, Watanabe T, et al. Purification and characterization of extracellular alkaline serine protease from *Stenotrophomonas maltophilia* strain S-1. Lett Appl Microbiol. 2005;41:253–257.

[55] Bhowmik T, Marth EH. Protease and peptidase activity of *Micrococcus* species. J Dairy Sci. 1988;71:2358–2365.

[56] Rodarte MP, Dias DR, Vilela DM, Schwan RF. Proteolytic activities of bacteria, yeasts and filamentous fungi isolated from coffee fruit (*Coffea arabica* L.). Acta Sci Agron. 2011;33:457–464.

[57] Bereded N, Curto M, Domig K, Beneberu F, Fanta S, Waidbacher H, et al. Metabarcoding analyses of gut microbiota of Nile tilapia (*Oreochromis niloticus*) from Lake Awassa and Lake Chamo, Ethiopia. Microorg. 2020;8:1040.

[58] Jaouadi NZ, Rekik H, Badis A, Trabelsi S, Belhoul M, Yahiaoui AB, et al. Biochemical and molecular characterization of a serine keratinase from *Brevibacillus brevis* US575 with promising keratin-biodegradation and hide-dehairing activities. Plos ONE. 2013;8:e76722.

[59] Li H-J, Tang B-L, Shao X, Lieu B-X, Zheng X-Y, Han X-X, et al. Characterization of a new S8 serine protease from marine sedimentary photobacterium sp. a5–7 and the function of its protease-associated domain. Front Microbiol. 2016;7:02016.

[60] Saggu SK, Jha G, Mishra PC. Enzymatic degradation of biofilm by metalloprotease from *Microbacterium* sp. SKS10. Front in Bioeng Biotechnol. 2019;7:192.

[61] Zhou C, Qin H, Chen X, Zhang Y, Xue Y, Ma Y. A novel alkaline protease from alkaliphilic *Idiomarina* sp. C9-1 with potential application for eco-friendly enzymatic dehairing in the leather industry. Sci Rep. 2018;8:16467.

[62] Yildirim V, Baltaci MO, Ozgencli I, Sisecioglu M, Adiguzel A, Adiguzel G, et al. Purification and biochemical characterization of a novel thermostable serine alkaline protease from *Aeribacillus pallidus* C10: a potential additive for detergents. J Enzyme Inhib Med Chem. 2017;32:468–477.

[63] Chellappan S, Jasmin C, Basheer SM, Kishore A, Elyas KK, Bhat SG, et al. Characterization of an extracellular alkaline serine protease from marine *Engyodontium album* BTMFS10. J Ind Microbiol Biotechnol. 2011;38:743–752.

[64] Niyonzima FN, More SS. Purification and characterization of detergent-compatible protease from *Aspergillus terreus* gr. 3 Biotech. 2015;5:61–70.

