## Supplementary Figure S1 for "Substrate preferences, phylogenetic and biochemical properties of proteolytic bacteria present in the digestive tract of Nile tilapia"

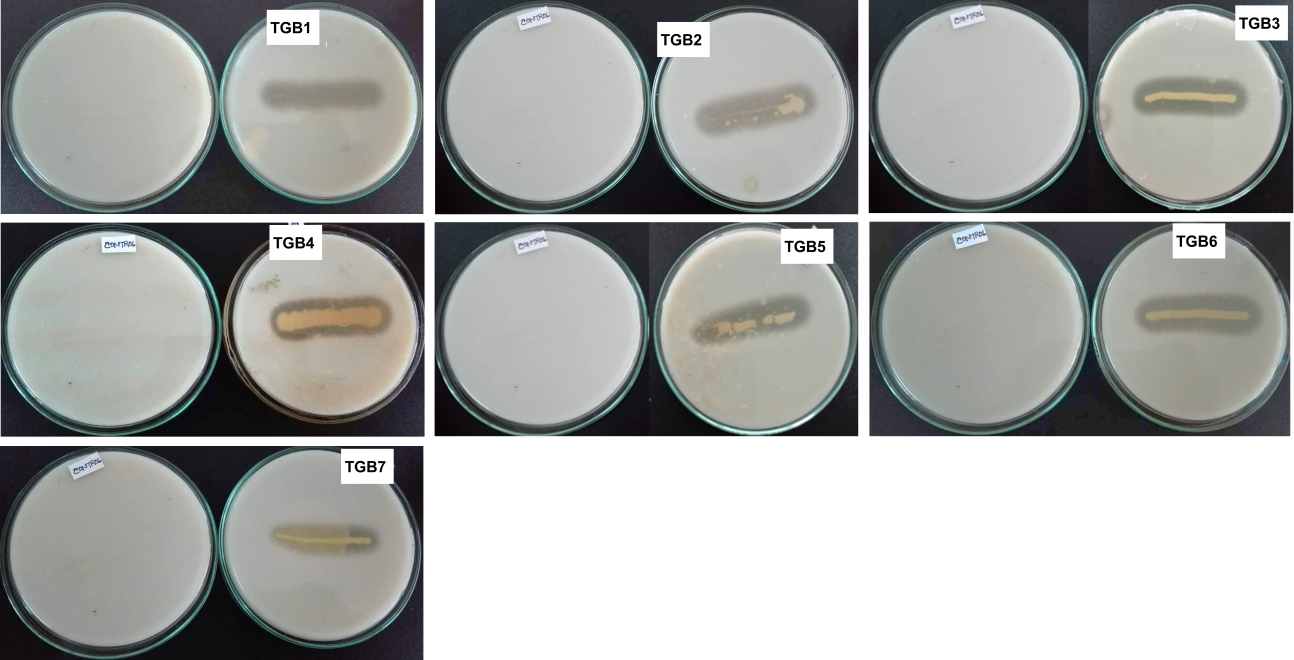

**Supplementary Figure S1.** Screening for proteolytic activity of the isolates, indicated by clear zones around the colonies grown on media containing casein. For each isolate, the plate at the left shows negative control in which bacteria without proteolytic activity was streaked on the media.
